# Nuclear transcriptomes of the seven neuronal cell types that constitute the *Drosophila* mushroom bodies

**DOI:** 10.1101/412569

**Authors:** Meng-Fu Maxwell Shih, Fred P. Davis, Gilbert Lee Henry, Josh Dubnau

**Author notes:** To whom correspondence should be addressed.; Correspondence may also be addressed to G.L. Henry. These authors contributed equally to this work (Meng-Fu Maxwell Shih, Fred P. Davis).

## Abstract

The insect mushroom body (MB) is a conserved brain structure that plays key roles in a diverse array of behaviors. The *Drosophila melanogaster* MB is the primary invertebrate model of neural circuits related to memory formation and storage, and its development, morphology, wiring, and function has been extensively studied. MBs consist of intrinsic Kenyon Cells that are divided into three major neuron classes (γ, α’/β’ and α/β) and 7 cell subtypes (γd, γm, α’/β’ap, α’/β’m, α/βp, α/βs and α/βc) based on their birth order, morphology, and connectivity. These subtypes play distinct roles in memory processing, however the underlying transcriptional differences are unknown. Here, we used RNA sequencing (RNA-seq) to profile the nuclear transcriptomes of each MB neuronal cell subtypes. We identified 350 MB class- or subtype-specific genes, including the widely used α/β class marker *Fas2* and the α’/β’ class marker *trio*. Immunostaining corroborates the RNA-seq measurements at the protein level for several cases. Importantly, our data provide a full accounting of the neurotransmitter receptors, transporters, neurotransmitter biosynthetic enzymes, neuropeptides, and neuropeptide receptors expressed within each of these cell types. This high-quality, cell type-level transcriptome catalog for the *Drosophila* MB provides a valuable resource for the fly neuroscience community.

## INTRODUCTION

*Drosophila melanogaster* is a powerful model system for behavioral neuroscience, which takes advantage of a relatively simple brain that expresses homologous genes, orchestrates a conserved yet highly diverse and elaborate suit of behaviors, during a short lifespan. Behavioral genetics in *Drosophila* affords the means to identify individual genes that function within identified neuronal cell types, whose connectivity and functional roles in behavior can be elucidated. The ability to form memories of past experience and to orchestrate adaptive and plastic changes in behavioral responses is an example of a fundamental field of behavioral neuroscience where *Drosophila* neurogenetics has made major contributions (1-4). Memory research in flies has led to the identification of fundamental cellular mechanisms of memory such as cAMP signaling and CREB-mediated gene transcription (1-5), and also has contributed to our understanding of how memories are processed in a complex neural circuit. A primary site of associative learning in insects is the mushroom body (MB) (1-4, 6-8), a paired brain structure that in *Drosophila* is comprised of approximately 2000 intrinsic Kenyon Cells (KCs) per hemisphere. MBs in fruit flies are critical sites of olfactory, visual and gustatory learning (1-4, 9, 10), and also play important roles in other behavioral contexts such as temperature preferences (11), sleep (12) and responses to ethanol exposure (13).

MB dependent plasticity is one of the most intensely studied aspects of invertebrate neurobiology. The morphology and developmental lineage of the neurons that populate the MB in *Drosophila*, as well as the identity and morphology of most of their neuronal inputs and outputs, have been fully characterized (14-17). Many functional manipulations of both neural activity and signaling pathways relevant to plasticity have been conducted within each of the identified neuronal cell types in this circuit (18-30). Functional imaging studies have established neural activity correlates in behaving animals (31). Together, these studies support the conclusion that the neurons of the MB play unique roles in memory acquisition, storage and retrieval. Moreover, memory storage over the course of minutes and hours after training relies on an evolving requirement for reverberating neural activity within a circuit that includes MB intrinsic neurons and the so-called extrinsic neurons that provide inputs and outputs (32, 33). In contrast to the increasingly deep understanding of the development, connectivity and functional requirements of each cell type in this circuit, there is a surprising paucity of data on differences in their transcriptional profiles.

MB KCs can be divided into three major classes, γ, α’/β’ and α/β, based on the projection patterns of the axons (34, 35). Extensive anatomical and functional characterization corroborates this classification as biologically relevant (1, 31, 32). Gene expression differences between these three classes of MB KCs have been studied using microarray (36), RNA sequencing (RNA-seq) (37) and single-cell RNA-seq (38, 39). However, it is known that the three KC classes can be further separated into seven subtypes: γd, γm, α’/β’ap, α’/β’m, α/βp, α/βs and α/βc KCs. The functional relevance of this further subdivision is supported by expression of split-GAL4 lines, analysis of the axonal projection patterns of individual neurons from each cell subtype (17, 40) and investigation of their functional roles in behavior (19, 41-43). Until now, there have been no attempts to identify the unique transcriptional programs that control/establish the identity of each of these seven cell subtypes, while cell clustering using a resource of enormous single-cell transcriptional profiles showed only three MB KC clusters (39). But the availability of intersectional genetics approaches that make use of split-Gal4 provide the means to investigate each of the subtypes individually (17, 40, 44).

To characterize the transcriptional programs of all seven MB neuronal subtypes, we combined genetic lines that specifically express in each (17) with tandem affinity purification of intact nuclei and RNA-sequencing (TAPIN-seq) (45) to profile their nuclear transcriptomes. These transcriptomes revealed ∽350 genes with either class-or subtype-specific expression, including several well-known and many new class or subtype markers. Moreover, our data provide a full accounting of the input-output signaling properties for each of these neuron subtypes including neurotransmitter biosynthetic machinery, neuropeptides and neurotransmitter and neuropeptide receptors.

## MATERIAL AND METHODS

### Fly stocks

Because quantitative traits such as gene expression profiles can be sensitive to genetic background, we created a newly isogenic subline of ***w***^*1118*^ (*isoCJ1*), which was itself derived from Canton-S wild type as an inbred line many years ago (46). To generate the isogenic strain, we used ten-generations of single male and female sibling crosses to generate 10 independent isogenic strains. MJ2 was selected based on its ability to form comparable olfactory short-term memory performance to the parental strain in the standard Pavlovian task (Figure S1). The nuclear envelope epitope tagged transgene *P{5XUAS-unc84∷2XGFP}attP40*, the pJFRC28 strain *P{10XUAS-IVS-GFP-p10}attP2* and each of the split-GAL4 inserts were backcrossed into this new MJ2 wild type strain for five generations to equilibrate each to this isogenic background. For each split-GAL4 combination, we separately backcrossed the GAL4 activating domain and DNA-binding domain components, and then combined the two hemi-drivers as a split-GAL4 line in the MJ2 background thereafter using standard balancer chromosomes that had themselves been equilibrated to the MJ2 strain. The *UAS-G2R-2 P{5XUAS-myr∷GFP-V5-p2A-His2B∷mCherry-HA}* reporter strain was generated using standard approaches. Flies were cultured on standard fly food at 25°C.

For imaging to characterize expression with each split-GAL4 strain that had been reconstituted in the MJ2 background, we used 2 – 5 day old male flies. For RNA-seq sample preparation, each split-GAL4 line was crossed to the *P{5XUAS-unc84∷2XGFP}attP40*. 2 – 5 day old adult progeny for each genotype were collected and frozen in liquid nitrogen.

### TAPIN purification of nuclei

Fly heads were first isolated with a customized sieve. 400 frozen heads were added to 6 mL of 20 mM sodium acetate pH 8.5, 2.5 mM MgCl_2_, 250 mM sucrose, 0.5% NP-40, 0.6 mM spermidine, 0.2 mM spermine, 1 mM DTT, 1× complete protease inhibitor (Sigma: 5056489001), 0.5 mg/mL torula RNA (ThermoFisher: AM7118), 0.6 mg/mL carboxyl coated Dynabeads (ThermoFisher: 14306D) and 1.6 mg anti-GFP antibody (ThermoFisher: G10362). Homogenization was carried out on ice by 50 tractions in a Dounce homogenizer using the tight pestle followed by filtration over either a 10 or 20 μm cup filter (Partec: 0400422314, 040042315).

Released chromatin and broken nuclei were adsorbed to carboxyl coated magnetic beads for 30 minutes at 4°C with constant rotation. Unbound antibody was removed by incubating the sample on ice for 20 minutes with 100 mL of washed UNOsphere SUPra resin (Bio-Rad: 1560218). After the resin was removed on a 10 μm cup filter and the carboxyl beads on a magnet stand, the nuclei-containing supernatant was mixed with an equal volume of 500 mM sodium acetate pH 8.5, 250 mM sucrose, 6 mM EGTA, 6 mM EDTA, 0.6 mM spermidine, 0.2 mM spermidine, 1 mM DTT, 1× complete protease inhibitor, 0.25 mg/mL torula RNA and 30 mL Protein A Dynabeads (ThermoFisher: 10002D). A 2-hour incubation on ice with occasional agitation was used to recover tagged nuclei. Bead-bound nuclei were then recovered on a magnet stand and washed twice with 250 mM sodium acetate pH 8.5, 250 mM sucrose and 0.1% NP-40. Nuclei were then released at 37°C for 1 hour by incubation in 50 μL of 10 mM Tris pH 7.5, 2.5 mM MgCl_2_, 0.5 mM CaCl_2_, 250 mM sucrose, 0.1% NP-40, 1 mg/mL torula RNA, 40 units RNAsin (Promega: N2515), 2 units DNAseI (NEB: M0303L), 320 units IdeZ protease (NEB: P0770S). The sample was diluted to 100 μL with 10 mM Tris pH 7.5, 2.5 mM MgCl_2_, 0.5 mM CaCl_2_, 250 mM sucrose and 0.1% NP-40, EGTA was added to 1 mM and the suspension was rapidly triturated 100 times. After returning the sample to a magnet stand, 90 μL of buffer containing released nuclei was removed and added to 1.5 μL of Protein G Dynabeads (ThermoFisher: 10004D) that were previously resuspended in 10 μL of 10 mM Tris pH 7.5, 2.5 mM MgCl_2_, 0.5 mM CaCl_2_, 250 mM sucrose and 0.1% NP-40. The second binding reaction was run for 1 – 3 hours on ice with occasional agitation, followed by 2× 250 μL washes in 10 mM Tris pH 7.5, 2.5 mM MgCl_2_, 0.5 mM CaCl_2_, 250 mM sucrose and 0.1% NP-40. Prior to the last wash a 5 μL aliquot was removed for quantitation and the remainder of the sample was solubilized in Arcturus Picopure RNA extraction buffer (ThermoFisher: KIT0204).

### RNA-seq library construction

Nuclear RNA was DNAseI treated and purified using the Arcturus PicoPure system exactly as instructed by the supplier. Purified RNA was mixed with a 1:100,000 dilution of ERCC standard RNA (ThermoFisher: 4456740) and amplified using the Nugen Ovation v2 system (Nugen: 7102-32). cDNA was then blunted, ligated to barcoded linkers (Nugen: 0319-32, 0320-32) and sequenced in paired-end mode on an Illumina HiSeq2500 to 125 nt read lengths.

### RNA-seq data analysis

We trimmed RNA-seq reads (5nt from the 5’ end of the forward read, using seqtk option “trim-b 5”; https://github.com/lh3/seqtk) to remove non-transcript sequences introduced by the NuGen Ovation kit and then pseudo-aligned these to the Drosophila transcriptome (ENSEMBL release 91, BDGP6) using kallisto (47) to estimate transcript abundances. We also included sequences for the synthetic ERCC spike-in species and TAPIN reporter in the transcriptome index. After pseudo-alignment, we removed ERCC, TAPIN reporter, and ribosomal RNA entries and renormalized the transcript abundance matrix to units of transcripts per million (TPM). To visualize TAPIN-seq signal across the genome, we also aligned trimmed reads to the whole genome (BDGP6, dm6) using STAR (48), created bigWig genome tracks (deeptools) (49), and visualized them in the IGV genome browser (50).

To identify class- and subtype-enriched genes, we performed differential expression analysis using the estimated counts from kallisto as input to limma (51), voom (52), and quantile normalizing the expression levels to account for differences in the number of genes detected in each sample (Table S1). We used criteria of at least 10 TPM abundance in one sample, at least two-fold difference in expression, and 5% false discovery rate to identify differentially expressed genes.

### Immunohistochemistry

Immunohistochemistry was performed essentially as in a previous report (53). Brains were dissected in isotonic PBS and immediately transferred to 4% paraformaldehyde in PBS for a 30-min fixation at room temperature. Fixed brain samples, were rinsed with isotonic PBS and incubated in PBS containing 2% Triton X-100, 10% normal goat serum (NGS; Penetration & Blocking Buffer) while being subjected to a degassing procedure (54). Brain samples were agitated in the same buffer at 4°C overnight. Brains were then transferred to primary antibodies diluted with PBS containing 0.25% Triton X-100, 1% NGS (Dilution Buffer) and agitated at 4°C for 1–3 day. After primary antibody incubation, the brain samples were washed in PBS containing 1% Triton X-100, 3% NaCl (Washing Buffer) three times before they were moved to the 1:250 diluted secondary antibodies for one day agitation at 4°C. For the biotin amplification staining (Figure 4, S3 & S5A), samples were washed three times and agitated in the 1:500 diluted Alexa Fluor 635 streptavidin (Thermo Fisher Scientific, USA: S-32364) at 4°C for 1 day. Finally, the immunolabeled brain samples were washed three times, cleared and mounted in a drop of *FocusClear* (CelExplorer Labs, Taiwan: FC-101) between two coverslips separated by a spacer ring of ∽200 μm thickness, so the brain samples were not flattened. The Penetration & Blocking Buffer and Dilution Buffer contain additional 0.02% Sodium Azide as a preservative. For GAL4 line characterization, 1:100 dilution of mouse anti-dlg1 (4F3, deposited to the DSHB, USA by Goodman, C.) plus 1:250 dilution of rabbit anti-GFP (Thermo Fisher Scientific, USA: A-6455) primary antibody and 1:250 dilution of secondary antibody of Alexa Fluor 633-conjugated goat anti-mouse (Thermo Fisher Scientific, USA: A-21052) and Alexa Fluor 488-conjugated F(ab’)2- goat anti-rabbit (Thermo Fisher Scientific, USA: A-11070) were used. For the MB marker gene confirmation, a 1:4000 dilution of rabbit anti-sNPFp (55) or 1:20 dilution of mouse anti-Fas2 (1D4, deposited to the DSHB, USA by Goodman, C.) or 1:20 dilution of mouse anti-trio (9.4A, deposited to the DSHB, USA by Hama, C.) primary antibody, 1:250 dilution of secondary antibody of biotin-conjugated goat anti-rabbit (Thermo Fisher Scientific, USA: 65-6140) or biotin-conjugated goat anti-mouse (Thermo Fisher Scientific, USA: D-20691) were used. For the GABAergic identification staining, a 1:250 dilution of mouse anti-GFP (MilliporeSigma, USA: 11814460001) together with 1:250 dilution of rabbit anti-GABA (MilliporeSigma, USA: A2052) or 1:500 dilution of rabbit anti-Gad1 (56) or 1:400 dilution of rabbit anti-VGAT (57) primary antibodies, and 1:250 dilution of secondary antibody of Alexa Fluor 488-conjugated goat anti-mouse (Thermo Fisher Scientific, USA: A-11029) together with biotin-conjugated goat anti-rabbit or Alexa Fluor 647-conjugated goat anti-rabbit (Thermo Fisher Scientific, USA: A-21244) were used.

**Figure 4.**
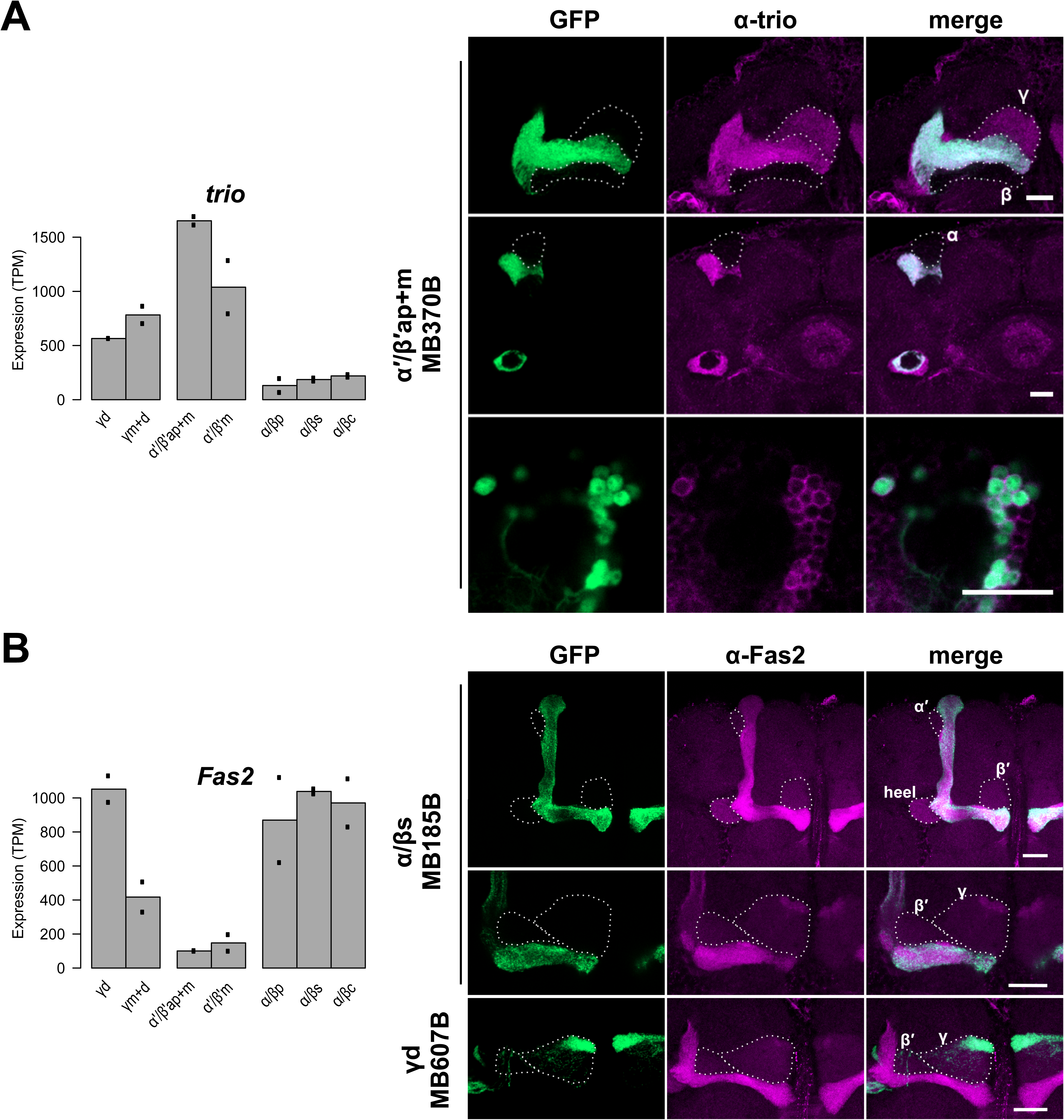
(A) *trio* is depleted in α/β KCs. Whole mount anti-trio immunostaining confirmed strong signal in the MB α’/β’ lobes, moderate signal in the γ lobe, and no signal in the α/β lobes. The cell bodies of MB α’/β’ KC class also showed immunoreactivity. (B) *Fas2* is depleted in MB α’/β’ KC class. Whole mount anti-Fas2 immunostaining confirmed stronger signal in the MB α/β and γd lobes. (C) *sNPF* is depleted in MB α’/β’ KC class. Whole mount anti-sNPF precursor immunostaining confirmed no detectable signal in the MB α’/β’ lobes. Among the immunoreactive α/β and γ lobes, the α/βp lobes showed the strongest signal. In each plot the bars represent the mean TPM, and the dots represent individual replicate values. Scale bars represent 20 μm. Expression patterns of the split-GAL4 lines were reported by *P{10XUAS-IVS-GFP-p10}attP2*.

### Confocal imaging and post-imaging processing

Brain samples were imaged under a Zeiss LSM 800 confocal microscope with a 40X C-Apochromat water-immersion objective lens (N.A. value 1.2). The settings for scanning were manually adjusted. To overcome the limited field of view when imaging the GAL4 expression patterns, we scanned each brain in two parallel stacks of confocal images with some overlap between the two brain hemispheres, with a voxel size of 0.31 X 0.31 X 1.25 μm. We then stitched the two image stacks into a single data set with ‘Pairwise stitching’ function in Fiji (58, 59), segmented the brain region based on the dlg1 staining channels with 3D Slicer (https://www.slicer.org) (60), and made a ‘Z projection’ with Fiji (59). The MB subtype models were constructed from the GFP channel of the confocal images used for projections, by using the 3D Slicer to segment, show 3D, and conduct smoothing.

### Cell counting

The confocal images for cell counting were acquired with a voxel size of 0.16 X 0.16 X 1.00 μm. The GFP channel was first used to identify KCs, and then the ‘Cell Counter’ plugin in Fiji was used to count all detectable nuclei in the mCherry channel (58). For each line, we counted three hemispheres from three different animals.

## RESULTS

### TAPIN-seq profiling of MB neuronal cell subtypes

To label MB subtypes, we used seven split-GAL4 lines: MB607B (γd), MB131B (γm+d), MB370B (α’/β’ap+m), MB418B (α’/β’m), MB371B (α/βp), MB185B (α/βs) and MB594B (α/βc) (17). We first backcrossed all the split Gal4 hemi-drivers and the nuclear-tag reporter (61) into MJ2, an isogenic Canton-S derivative (Methods). Because of the change in genetic background, we re-characterized the expression pattern of each split-GAL4 combination to confirm that the expected cell subtype-specific pattern had not been altered. We used a novel membrane-GFP-P2A-nuclear-mCherry dual label reporter (G2R for short; see Methods), which expresses membrane-tethered GFP and nuclear mCherry from a single transcript via a viral ribosome skip peptide (P2A) (62). The GFP label revealed neuronal morphology, thereby confirming MB subtype specificity, and the nuclear-mCherry marked each nucleus, which permitted an accurate cell count. Using G2R labeling, we found that each split-GAL4 combination yielded limited expression in only small numbers of neurons outside the MB, but exhibited strong expression in the annotated MB cell subtype (Figures 1A – 1G, 1A’ – 1G’ & Supplementary Movies 1 – 7) (17). Using high-resolution imaging (Figures 1A“–1G” & Supplementary Movies 8 – 14), we were able to count the total number of MB KCs of each subtype that is labeled by a given spilt-GAL4 combination (Table 1). The numbers of labeled cells for each MB KC subtype are in general agreement with a previous report (17) and it appears that most if not all of the neurons of each subtype are labeled with these Split-Gal4 combinations. Together, these results confirm the previously reported specificity and comprehensiveness of these split-GAL4 lines to label each of the MB KC neuronal subtypes.

**Table 1.** List of split-GAL4 lines labeling MB subtype (n=3)

| Driver | MB subtype | Number of cells |
| --- | --- | --- |
| MB607B | γd | 132.00 ± 6.03 |
| MB131B | γm+d | 574.33 ± 2.73 |
| MB370B | $\alpha'/\beta'$ap+m | 405.33 ± 18.80 |
| MB418B | $\alpha'/\beta'$m | 152.67 ± 3.71 |
| MB371B | $\alpha/\beta$p | 77.33 ± 2.85 |
| MB185B | $\alpha/\beta$s | 294.33 ± 16.68 |
| MB594B | $\alpha/\beta$c | 505.33 ± 9.67 |

**Figure 1.**
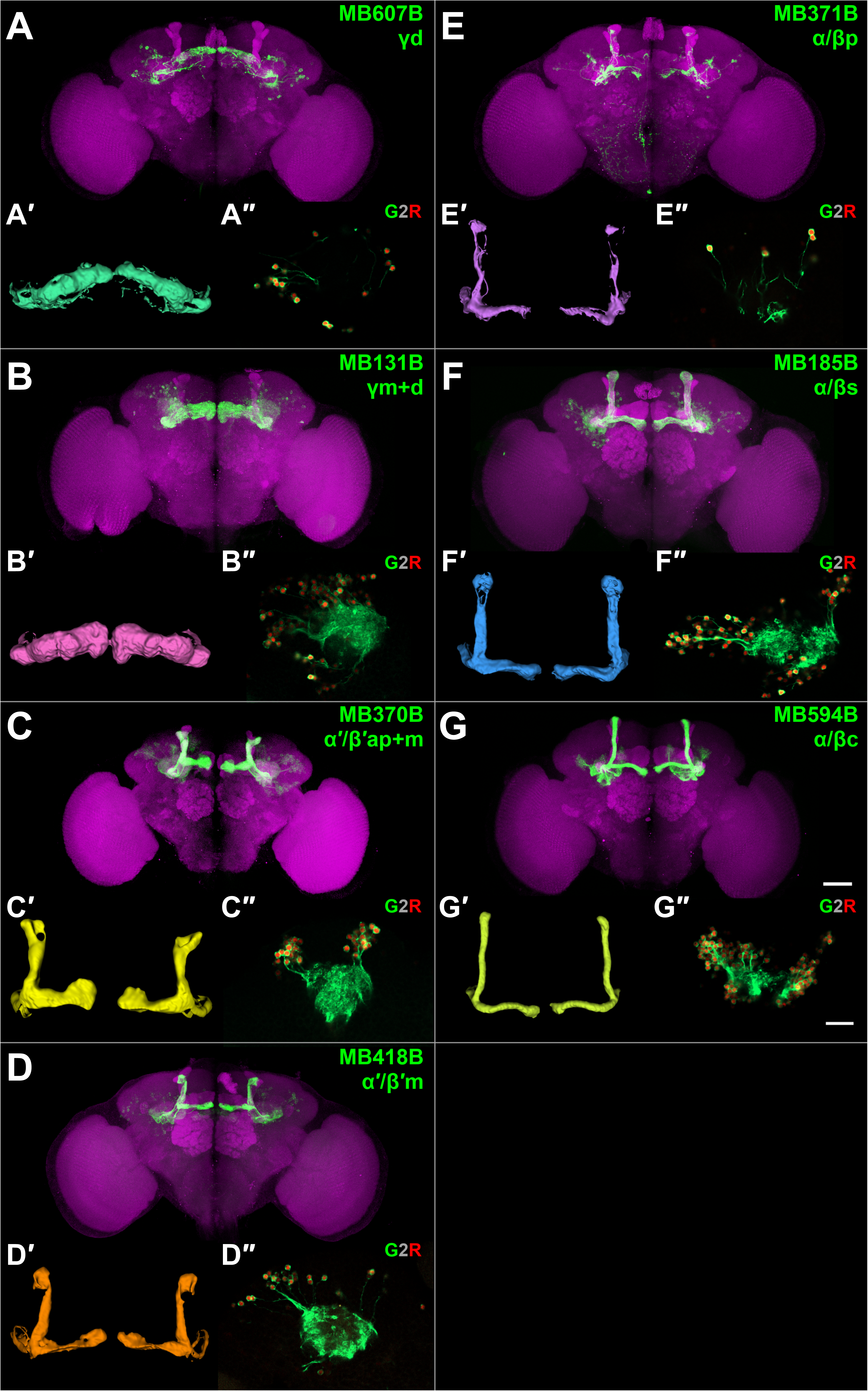
Characterizing mushroom body subtype drivers. (A-G) Expression pattern of the split-GAL4 driver lines used in this study. Green, the GFP plus anti-GFP immunoreactive signal; magenta, anti-dlg1 immunoreactive signal as a counterstain. The scale bar represents 50μm. (A’-G’) 3D model of the split-GAL4 expression pattern in MB. (A”-G”) Example of the high-resolution membrane-GFP- P2A-nuclear-mCherry dual label reporter (G2R for short; see Methods) images used for cell counting. The scale bar represents 20μm. Genotype: MB607B>G2R, *R19B03-p65.AD/UAS-G2R-2; R39A11- GAL4.DBD/+*, MB131B>G2R, *R13F02-p65.AD/UAS-G2R-2; R89B01-GAL4.DBD/+*, MB370B>G2R, *R13F02-p65.AD/UAS-G2R-2; R41C07-GAL4.DBD/+*, MB418B>G2R, *R26E07-p65.AD/UAS-G2R-2; R30F02-GAL4.DBD/+*, MB371B>G2R, *R13F02-p65.AD/UAS-G2R-2; R85D07-GAL4.DBD/+*, MB185B>G2R, *R52H09-p65.AD/UAS-G2R-2; R18F09-GAL4.DBD/+*, and MB594B>G2R, *R13F02- p65.AD/UAS-G2R-2; R58F02-GAL4.DBD/+*.

To profile the nuclear transcriptome of each MB subtype, we used TAPIN-seq (45), a modification of INTACT (61) that yields improved selectivity by use of a two-step purification (Figure S2A). The 7 characterized split-GAL4 combinations were used to express the nuclear membrane protein UNC84 fused with 2 copies of GFP (Figure 2A). For each split-Gal4 combination, tagged nuclei of a given MB KC subtype were purified from ∽400 fly heads. Nuclear RNA was then extracted and used to generate RNA-seq libraries (Figure S2B). We generated libraries from two independent biological replicates for each MB KC subtype, and paired-end sequenced them (Table S1). We estimated transcript abundances in each library using kallisto (47)(see Methods). The sequencing reads and estimated abundances are available in NCBI GEO (GSE119629; reviewer token wzczwumwlfitrmt).

**Figure 2.**
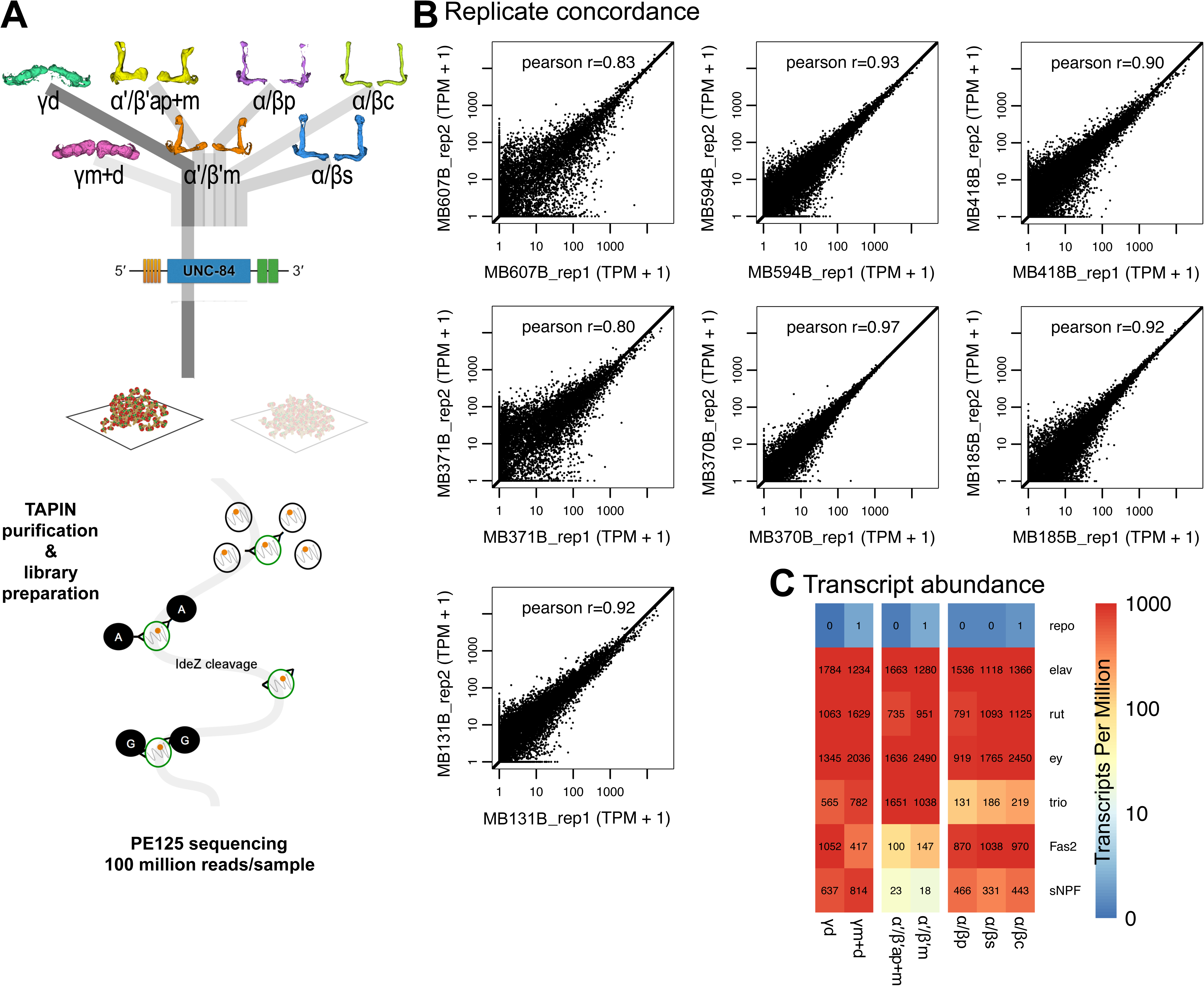
TAPIN-seq profiling of MB subtypes. (A) Driver lines expressing in the seven Kenyon cell subtypes were crossed with the TAPIN-seq reporter, which labels the nuclei in each subtype. Nuclear RNA from each subtype was used to generate RNA-seq libraries, which were sequenced in paired-end mode. (B) We estimated reproducibility of the TAPIN-seq measurements by calculating the Pearson correlation between estimated transcript abundances (log2 transformed Transcripts Per Million + 1). (C) The TAPIN-seq transcriptomes recover the neuronal marker *elav*, while not detecting the glial marker *repo*. The transcriptomes also recover the expected expression patterns of the known pan-Kenyon cell markers *ey* and *rut* as well as class-enriched genes *trio*, *Fas2*, and *sNPF*.

Transcript abundances were well correlated between replicate libraries (Figure 2B). We observed strong expression of the neuron-specific marker *elav* in all subtypes (1,118 – 1,784 TPM), contrasting with low levels of the glial-specific gene *repo* (0.1 – 1.3 TPM; Figure 2C), consistent with a high-fidelity purification of TAPIN labeled nuclei. We also detected strong and broad expression of genes expected in all MB neuron subtypes, such as the transcription factor *ey* (919 – 2,490 TPM) and components of the cAMP signaling pathway such as *rutabaga* (735 – 1,629 TPM) which encodes the Ca^2+^/calmodulin-activated adenylyl cyclase (Figure 2C) (34). The TAPIN-seq profiles also recovered the expected pattern of genes known to be enriched in individual classes, including *trio*, *Fas2*, and *sNPF*.

### Genes enriched in MB neuronal classes and subtypes

We next identified transcripts that are differentially expressed across MB neuron classes or subtypes using three criteria (Methods). We found 341 transcripts that are enriched or depleted in one of the three MB neuron classes (Figure 3A) and 57 that are enriched or depleted in one of the seven MB cell subtypes (Figure 3B; Table 2 for summary and Table S2 for the full gene list). To evaluate the accuracy of our differential expression analysis, we examined several genes encoding proteins reported to differentially label MB cell classes. Antibodies against trio, for example, are reported to label α’/β’ and γ classes (63). Indeed, the *trio* gene is identified in our analysis as an MB class-specific gene depleted in α/β KCs (average 179 TPM in α/β vs 1,345 TPM α’/β’ and 674 TPM γ), and we confirm that anti-trio immunoreactivity is localized in the MB α’/β’ and γ lobes, and their cell bodies (Figure 4A). Fas2 has been reported to exhibit strong immunoreactive signal in the MB α/β lobe class, weak signal in the γ lobe class and no signal in the α’/β’ lobe class (34). Consistent with this protein distribution, we identify *Fas2* transcripts as an MB class-specific gene depleted in α’/β’ KCs (124 TPM in TPM α’/β’ vs 959 TPM α/β and 734 TPM γ). At the cell type level, *Fas2* was enriched in in the γd subtype relative to γm+d (1,052 vs 417 TPM, respectively). Using immunolabeling, we confirmed this pattern of anti-Fas2 immunoreactivity. As previously reported, we detect strong immunoreactive signal of Fas2 in the α/β lobe class of neurons, somewhat weaker signal in the γ lobe class and no signal in the α’/β’ lobe class neurons. We further demonstrate that Fas2 immunoreactivity appears weaker in the γm subtype compared to γd subtype KCs (Figure 4B), which mirrors the prediction from the TAPIN-seq described above.

**Table 2.**
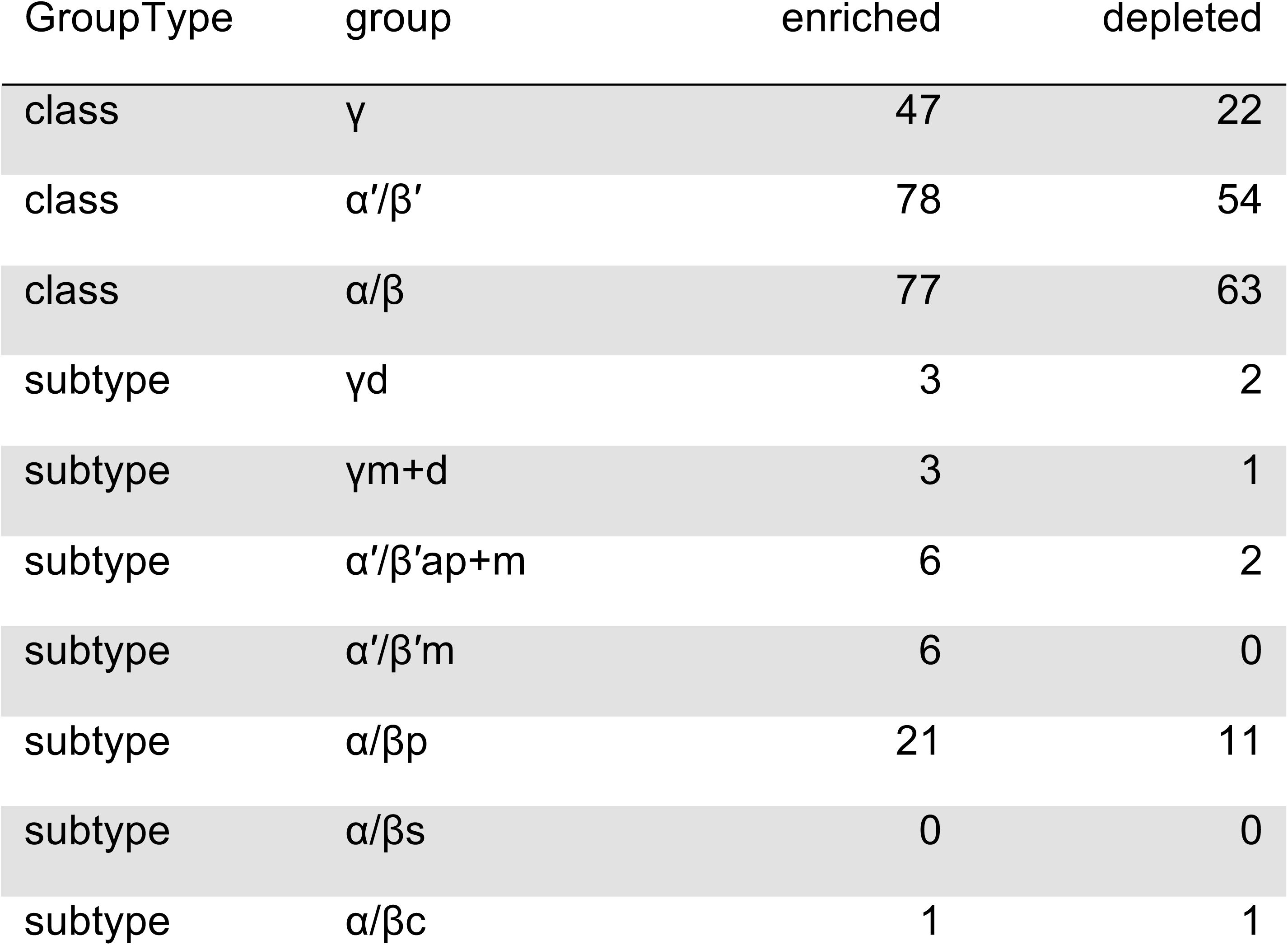
Number of genes enriched and depleted in individual MB classes and subtypes. Genes identified by limma/voom (q-value < 0.05, |fold change| > 2x).

**Figure 3.**
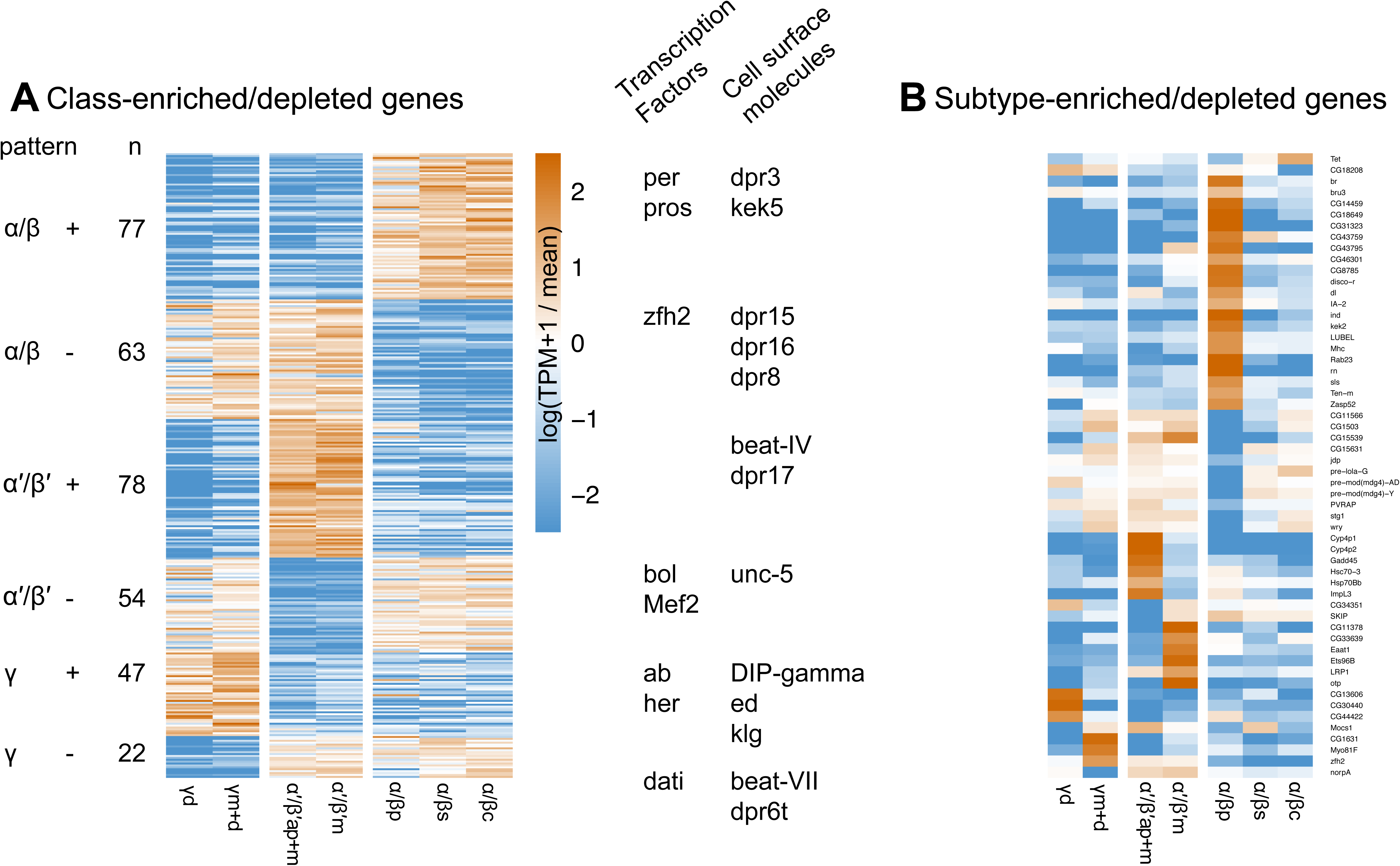
Differential expression of MB class-specific genes (A) and subtype-specific genes (B). Specific examples of up and downregulated genes for each subtype are indicated in (A).

We also examined the expression of the neuropeptide gene *sNPF* which we identified as an MB class-specific gene depleted in α’/β’ KCs (20 TPM in α’/β’ vs 414 TPM α/β and 725 TPM γ).Immunolabeling with an antibody against the sNPF precursor confirmed a previous report and is consistent with the TAPIN-seq results. We observe no detectable signal in α’/β’ KC class neurons and strong signal in α/β and γ classes (Figure 4C) (55). No noticeable signal was detected in the cell bodies of KCs (Figure S3), consistent with a previous description (55). At the protein level, we further noted elevated immunoreactivity in the MB α/βp subtype, which was not reflected by the TAPIN-seq results (Figure 4C). This discrepancy points either to limitations of TAPIN mediated profiling with these split-Gal4 lines and/or the importance of post-transcriptional regulatory mechanisms in determining the accumulation of the neuropeptide. Taken together, the identification of known differentially expressed transcripts and the immunostaining data for three identified examples broadly corroborate the fidelity of the TAPIN-seq results.

The differentially expressed genes also include several transcriptional regulators with class- and subtype-specific patterns (Figure 3). For example, we identify transcriptional regulators enriched in the γm+d (*zfh2*), α’/β’m (*Ets96b*, *otp*), α/βp (*br*, *bru-3*, *dl*, *disco-r*, *ind*, *rn*), and α/βc (*Tet*) subtypes. Although the abundance and significance of DNA methylation in *Drosophila* is unclear, enrichment of the *Tet* DNA methyltransferase is intriguing given the role of DNA methylation in memory formation in other insects (64) and mammals (65). The subtype-enriched expression could reflect a remnant of a functional expression pattern from an ancestral species with DNA methylation.

Several genes encoding cell-surface molecules were also differentially expressed across the subtypes. These genes included members of gene families previously implicated in specifying synaptic connectivity, including the defective proboscis extension response (Dpr) as well as the Dpr-interacting protein (DIP). Interactions between proteins from these families have been previously documented and shown to underlie synaptic connectivity in circuits including the visual system (66, 67). The expression patterns we observed for several cell-surface molecules suggest they might be involved in specifying class- and subtype-specific connectivity.

### Neurotransmitter output

To explore the functional utility of our TAPIN-seq measurements, we next focused on genes related to the input and output properties of the MB cell subtypes. Specifically, we examined genes encoding neurotransmitter biosynthetic enzymes, neurotransmitter transporters (Figure 5A), neurotransmitter receptors (Figure 5B), neuropeptides (Figure 5C), neuropeptide receptors (Figure 5D), and gap junction components (Figure 5E). A recent report has established that the primary neurotransmitter in the MB is acetylcholine (68). Because our method profiled each of the seven MB neuronal subtypes, we re-assessed this conclusion at a higher resolution. We confirmed that all seven MB cell subtypes express high levels of both *choline acetyltransferase* (*ChAT*; 75 – 198 TPM) and *vesicular acetylcholine transporter* (*VAChT*; 257 – 517 TPM).

**Figure 5.**
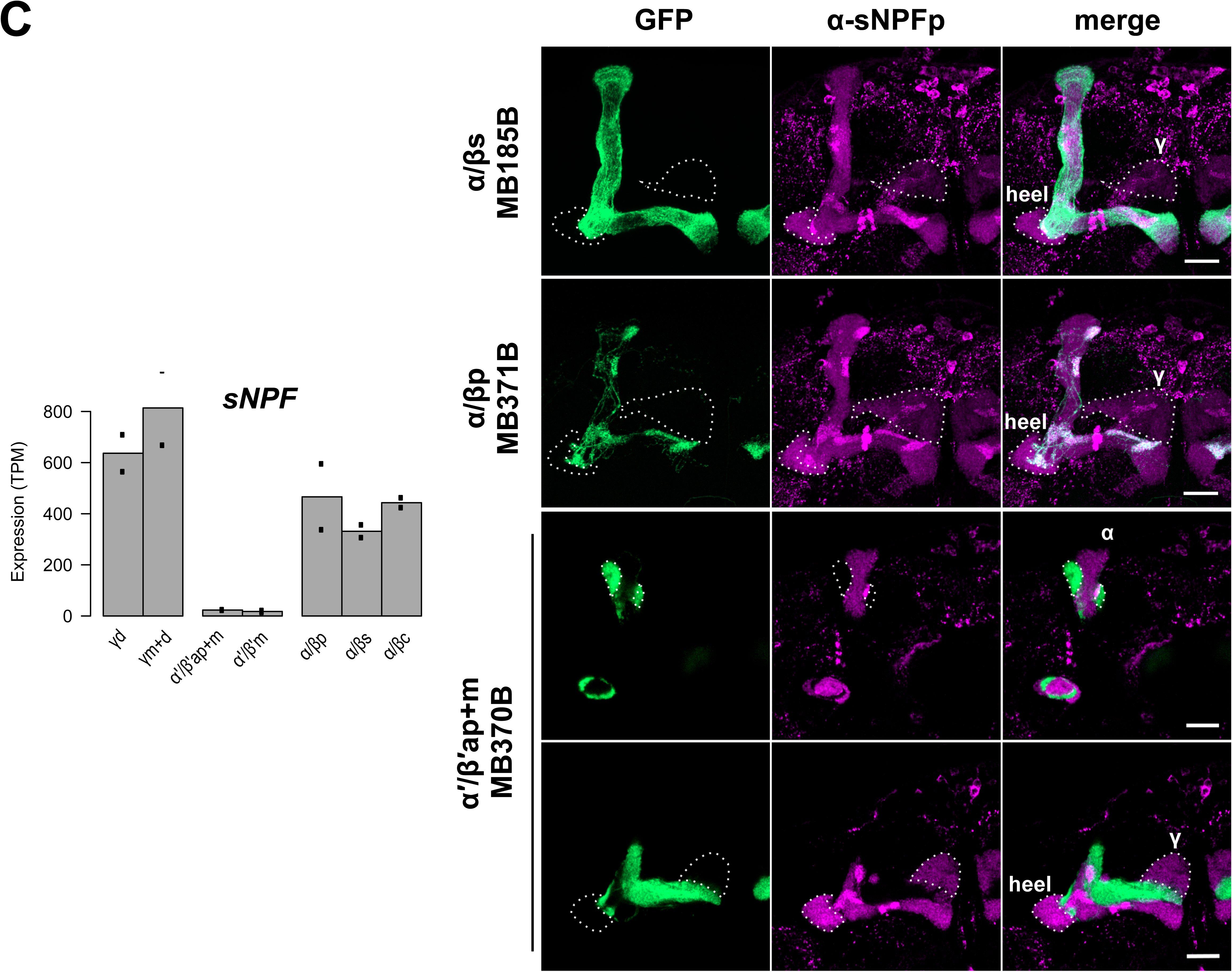
TAPIN-seq profiles of genes related to neurotransmitter biosynthetic enzymes, neurotransmitter transporters (A), neurotransmitter receptors (B), neuropeptides (C), neuropeptide receptors (D), and functional components of gap junctions (E).

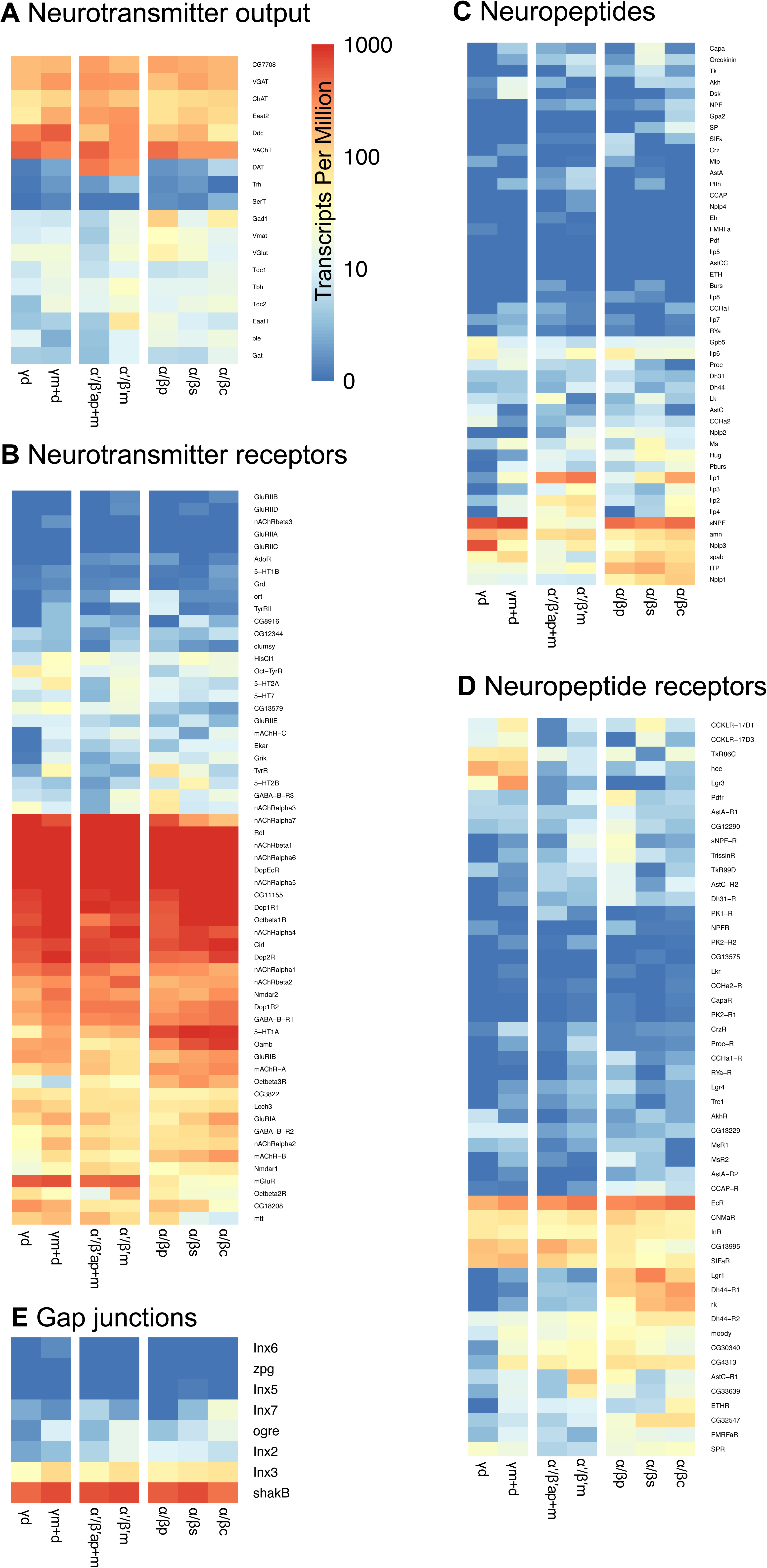

We next asked whether one or more KC subtypes might co-release other small molecule neurotransmitters, but found no strong argument to support such a conclusion. In a few cases, we see moderate expression of biosynthetic enzymes for other neurotransmitters including GABA, serotonin and dopamine. But in each case, there are findings that undermine the conclusion that these neurotransmitters are consistently produced/released by any of the 7 KC subtype. For example, we do see strong expression in all MB cell subtypes of *vesicular GABA transporter* (*VGAT;* 152 – 279 TPM), which is an essential transporter that is responsible for packaging the neurotransmitter GABA into synaptic vesicles (57). On the other hand, the gene for GABA biosynthetic enzyme, *Gad1*, is moderately expressed in only the α/βp (121 TPM) and α/βc subtypes of KCs (54 TPM) and at lower levels in the remaining cell subtypes (4 – 14 TPM). In principle, moderate *Gad1* expression might be consistent with the hypothesis that GABA is released from a fraction of α/βp and/or α/βc subtype neurons. To examine this possibility, we conducted immunofluorescence experiments with antibodies against GABA, Gad1, and VGAT but found no marked immunoreactivity in any α/βp KC subtype cell bodies (Figure S4). This result suggests that the most likely explanation for our observed *Gad1* pattern is that the split-GAL4 lines we used for α/βp and α/βc KC subtypes drive low levels of expression in some subset of GABAergic neurons outside the MB (Figure S4A).

A similar set of findings are apparent with Dopa decarboxylase (Ddc), a commonly used marker for dopaminergic or serotonergic neurons. Ddc catalyzes the decarboxylation of dopa to dopamine and 5- hydroxytryptophan to serotonin but not tyrosine to tyramine (69). We see fairly strong *Ddc* expression in all MB cell types (55 – 574 TPM), especially the γm+d (574 TPM) and γd (339 TPM) KCs. However, we do not see expression of *Tryptophan hydroxylase* (*Trh*; 0 – 3 TPM), which provides the first and rate-limiting step in the synthesis of serotonin. Nor do we detect *serotonin transporter* (*SerT*; 0 – 2 TPM) (70) in any of the 7 cell subtypes (Figure 5A). *Pale* (*ple*), which encodes tyrosine hydroxylase (TH) for dopamine synthesis, also is not highly expressed in any MB cell subtype (2 – 16 TPM; Figure 5A). Thus, we conclude that all MB KC subtypes likely release acetylcholine (37, 68) and no other small molecule neurotransmitters.

### Neurotransmitter receptors

We next examined expression profiles of small molecule neurotransmitter receptors. Dopamine has been established as a key input to MB for many behaviors including aversive and appetitive olfactory learning (28, 71-74), modulation of motivational state (75), regulated forgetting (76, 77), sleep (78), courtship behaviors (79, 80), and temperature preference (81). A network of dopaminergic neurons innervates all MB cell subtypes (17), and all four dopamine receptors, Dop1R1, Dop1R2, Dop2R and DopEcR, are expressed in all 7 MB KC subtypes (594 – 1152, 222 – 408, 444 – 1059, 3284 – 8335 TPM, respectively; Figure 5B) (28, 71, 82, 83; cf 84). We confirmed that the *Dopamine transporter* (*DAT)* is preferentially expressed in both MB α’/β’ap and α’/β’m cell subtypes (397 and 262 TPM in α’/β’ and α’/β’m, respectively, vs 0.2 – 4.3 TPM in other subtypes; Figure 5A) (38, 39), suggesting that precise temporal control of dopaminergic signaling may be at play in α’/β’ KCs. We also observe equivalently high levels of expression in all MB KC subtypes for all six GABA receptor genes. Among the three GABA_A_ receptors, *Resistance to dieldrin* (*Rdl*) is strongly expressed (5,021 – 9,548 TPM), consistent with previous findings (85, 86), while *Ligand-gated chloride channel homologue 3* (*Lcch3*) is moderately expressed (70 – 153 TPM), and *Drosophila Glycine receptor* (*Grd*) is not detected (0.1– 0.8 TPM). All the GABA_B_ receptors are broadly expressed in MB except for *GABA-B-R3* (1 – 61 TPM; Figure 5B). These findings are consistent with a report that establishes the importance of GABAergic feedback from the anterior paired lateral neurons to MB KCs (87), mediated by both ionotropic GABA_A_ and metabotropic GABA_B_ receptors (88).

Serotonergic signaling in the MB also is involved in olfactory memory formation, sleep regulation and stress response modulation (89-92). We found that among five serotonin receptors, only *5-HT1A* is expressed in the MB, with an enrichment in the α/β lobe KC class (Figure 5B). This is consistent with the finding that dorsal paired medial (DPM) neurons release serotonin onto 5-HT1A receptors expressed in α/β KCs to support anesthesia-resistant memory formation (89). A previous report observed *5-HT1B-GAL4* expression and 5-HT1B immunoreactivity in the γ KC class (93). We did not detect nuclear *5-HT1B* transcripts by TAPIN-seq (0.1 – 1.0 TPM), but we cannot rule out the possibility that higher levels of transcripts and protein are present in the cytoplasm, beyond detection in this nuclear transcriptome.

MB KCs receive nicotinic acetylcholine receptor (nAChR)-mediated synaptic transmission from antennal lobe projection neurons (94). We found the nAChR subunits *nAChRalpha1*, *nAChRalpha4*, *nAChRalpha5*, *nAChRalpha6*, *nAChRalpha7*, *nAChRbeta1* and *nAChRbeta2* are strongly expressed in all seven MB cell subtypes, but *nAChRalpha3* and *nAChRbeta3* transcripts are absent or undetectable (Figure 5B). Two muscarinic acetylcholine receptors (mAChRs), *mAChR-A* and *mAChR-B*, are also expressed in the MB with an enrichment in α/β lobe KC class (Figure 5B & Table S2).

We also examined the expression of the six known octopamine receptors (69). We found that *Oamb* and *CG18208* (recently characterized as *Octα2R*) (95), which are α-adrenergic-like receptors, are class- and subtype-specific genes, respectively (Table S2). *Oamb* TAPIN-seq signal is strongly detected in α/β class KCs and far lower levels are seen in α’/β’ class KCs (Figure 5B) (34, 96). The three β-adrenergic-like receptors — *Octβ1R*, *Octβ2R* and *Octβ3R* — are all detected in the MB with variable levels across cell subtypes (cf 53 for Octβ2R immunolabeling).

Although transient glutamate immunoreactivity has been shown in α/βc lobe KC class of young adult, *VGlut* expression has never been observed in the MB KCs (97, 98). Consistent with this conclusion, we observe low *VGlut* transcript abundance (5 – 50 TPM; Figure 5A). We also confirmed that the NMDA receptors, both *Nmdar1* and *Nmdar2*, are broadly expressed in the MB (Figure 5B) (99, 100); ionotropic receptors *GluRIA* and *GluRIB* are also expressed (Figure 5B). Flies also have a unique metabotropic glutamate receptor called mGluR, which has been previously observed by immunolabeling throughout the adult brain, but minimally in the MB lobes (101). Here, with the cell type resolution of our dataset, we identified *mGluR* as an MB class-specific gene that is expressed in a subset of KCs — *mGluR* is depleted in α/β class KCs (40 TPM; Table S2) relative to both α’/β’ and γ classes of KCs (504 and 612 TPM, respectively; Figure 5B).

### Neuropeptides

In addition to small neurotransmitter systems, we also examined expression of neuropeptides and neuropeptide receptors. We consistently detect expression of a small group of well-characterized and putative neuropeptides. Unexpectedly, this includes *amnesiac* (*amn*), which is detected at fairly robust levels in all 7 MB KC subtypes (52 – 175 TPM) — although this small gene resides in the intron of another gene, *Hers*, which is highly expressed (218 – 239 TPM), thus complicating the accurate estimation of *amn* abundance. Previous work has established a requirement during memory formation for *amn* expression in a single pair of DPM neurons outside MB (102). *sNPF* expression appears as a class-specific transcript, with high levels in α/β and γ KC classes and very little expression in the α’/β’ KC class (414, 725, 20 TPM, respectively; Figure 4C). This is consistent with previous reports (55) and we further confirm this expression pattern by immunostaining (Figure 4C). Another notable class-specific neuropeptide is *Drosophila Insulin-like peptide 1* (*Ilp1*) (103), expressed at high levels in α’/β’ class of KCs (284, 382 TPM in α’/β’, α’/β’m, respectively) and α/βc subtype (220 TPM) but not in most other MB KC subtypes (0.8 – 55 TPM; Figure 5C). As with neuropeptides, we also detect a panel of neuropeptide receptors, some of which are robustly expressed in all MB KC subtypes (eg, *Ecdysone receptor*), some of which are class specific (eg, *Dh44-R1* and *hector*), and some of which are enriched or depleted in one or more subtypes of neurons (eg, *AstC-R1* and *ETHR*; Figure 5D).

Finally, because of a report that gap junctions may form between MB KCs of different classes and play a role in visual learning (104), we examined the expression levels of the eight gap junction genes in MB cell subtypes. *shakB* is strongly expressed (433 – 751 TPM), and *Inx3* moderately (29 – 107 TPM), in all MB cell subtypes (Figure 5E). Although *Inx5* and *Inx6* are reportedly required in the α/β and α’/β’ KC classes for visual learning and memory (104), we do not detect either gene in any of the 7 MB KC subtypes (0 – 0.2 TPM, 0 – 0.5 TPM, respectively; Figure 5E). We cannot rule out the possibility that functionally relevant levels of expression are below our detection limit or that cytoplasmic RNA levels are higher.

## DISCUSSION

Our results establish a high-quality, neuronal cell type-level transcriptome for *Drosophila* MB. We identified 350 differentially expressed genes, which includes most of the previously reported MB lobe (class specific) markers and many novel class-specific or cell subtype-specific profiles of expression. In addition to the subtype level resolution of our experimental design, the TAPIN approach that we used also offers several advantages and technical differences with these prior approaches. First, because TAPIN is compatible with flash frozen tissue as the input, the method introduces minimal disturbance to the endogenous transcriptome as compared to more lengthy procedures for purification of neurons for expression profiling. Second, it may be relevant that TAPIN explicitly profiles nuclear RNAs, likely enriching for actively transcribed/nascent transcripts versus abundant ones that are stably maintained in the cytoplasm. Thus, it would be attractive to apply this method to profile transcriptional response to behavioral perturbations.

Several previous studies have used genome-wide methods to profile expression in the *Drosophila* MB (36-39, 105). Perrat et al. used a microarray-based approach to profile expression of each of the three major classes of MB KCs (purified by flow cytometry from dissociated brains) and compared these profiles with expression in the rest of the brain. They first focused on the expression of transposons (36), and subsequently used the same transcriptome dataset to discover that MB KCs are cholinergic (68) based on expression of biosynthetic enzymes. Crocker et al. used an RNA-seq-based approach to profile expression in relatively small pools of physically isolated α/β and γ class neurons to search for memory-related changes in gene expression (37). Most recently, Croset et al. and Davies et al. used droplet-based single cell sequencing to profile the *Drosophila* brain, and by clustering the single cells they were able to identify the three MB classes, but not the further sub-division into neuronal subtypes (38, 39).

Although we used a different profiling method and resolved transcriptomes at the cell subtype rather than class level, our findings are broadly compatible with prior reports (36-39, 68). Our dataset reveals strong expression of both *ChAT* and *VAChT*, consistent with the conclusion that MBs are cholinergic (68). Our findings further support the conclusion that all of the individual MB KC subtypes are cholinergic. The datasets also are consistent in the expression of known class-specific markers. One notable difference is that Crocker et al. reported high levels of expression of the *5-HT1B* receptor in both α/β and γ classes of KCs (37), and Davie et al. also observed 5-HT1B expression in α/β and γ KCs single cell clusters (39). In contrast, we see no evidence for expression of this receptor in our TAPIN-seq profiles (0.1 – 1 TPM). This difference could reflect methodology: Crocker and Davie both measured 5-HT1B receptor transcripts in the cytoplasm while we measured the levels that are actively transcribed or present in the nucleus. This technical difference could be especially relevant for neurotransmitter receptors, some of which can be translated locally at dendrites (106).

Our dataset is the first to profile expression in this brain region at neuronal subtype resolution. This level of resolution is critical given the wealth of data on the functional differences of each MB KC subtype in *Drosophila* behaviors. Our dataset provides a full accounting within each of the MB KC subtypes of the profiles of expression of the cellular machinery to produce and receive neurotransmission, including small molecule transmitters and their receptors, neuropeptides and neuropeptide receptors and subunits of gap junctions. It is noteworthy that the TAPIN expression dataset supports the conclusions that all the adult MB KC subtypes are cholinergic, and that none of the subtypes express genes that would suggest the co-release of GABA, dopamine, glutamate, or serotonin. On the other hand, we detect expression of a spectrum of neuropeptides and their receptors. This observation is consistent with the hypothesis that MB KCs may co-release both acetylcholine and several neuropeptides (107).

In addition to these findings with regards to the inputs and outputs, we identified 350 differentially expressed genes including many that distinguish MB KC classes or even individual cell subtypes. MB α/βp subtype showed 21 enriched genes and 11 depleted genes, contrasting with two other subtypes in the α/β class and two other classes. This uniqueness is supported by its unique odor responses (19) and connectivity (108). Despite the limitation in the methodology that we profile the MB γm subtype using the spilt-GAL4 line MB131B that has minor expression in γd, and profile the α’/β’ap subtype using MB370B that has minor expression in α’/β’m (Figure 1 & Table 1), we still identified distinct sets of enriched/depleted genes (Figure 3 & Table S2), indicating the differences between two subtypes in MB γ or α’/β’ classes.

Our dataset provides a valuable resource for the fly neuroscience community to conduct functional studies. For example, our data provide a list of previously unknown class specific and sub-type specific transcripts, whose impact on the functional differences between these neurons are not known (Figure 3; Table S2). An arsenal of genetic tools to manipulate any gene’s function within each of these cell subtypes already exists (109, 110). In addition to olfactory associative memory, MBs also play fundamental roles in other forms of memory including visual and gustatory (9, 10), temperature preference (11), courtship behaviors (79, 80), stress response (92), food-seeking (111), sleep (12) and responses to ethanol (13). This dataset will facilitate the discovery of neural mechanisms for each of these conserved behaviors.

## AVAILABILITY

All code used to analyze RNA-seq results and create the figures and tables in this manuscript are available in the GitHub repository (http://github.com/fredpdavis/mushroombody).

## ACCESSION NUMBERS

The sequencing reads and estimated abundances are available in NCBI GEO (GSE119629; reviewer token wzczwumwlfitrmt).

## SUPPLEMENTARY DATA

Supplementary Data are available at NAR online.

## ACKNOWLEDGEMENT

We thank Y. Aso and G. Rubin (Janelia Research Campus) for the split-GAL4 lines and pJFRC28 reporter, J. Veenstra (Université de Bordeaux 1, France) for the rabbit anti-sNPFp antibody, D. Krantz (University of California, Los Angeles) for the rabbit anti-VGAT antibody, F. Jackson (Tufts University) for the rabbit anti-Gad1 antibody and the Developmental Studies Hybridoma Bank (The University of Iowa), created by the NICHD, for the mouse monoclonal antibodies against dlg1, Fas2 and trio. We thank Y.-H. Chang for the G2R fly strain and W.-W. Liao (Washington University in St. Louis) for technical assistance in analysis of the alignment files. We also are grateful to J. Azpurua, J. Beshel, Y.-H. Chang, R. Keegan, S. Krupp, L. Talbot for helpful discussions or comments on the manuscript. This work utilized the computational resources of the NIH HPC Biowulf cluster (http://hpc.nih.gov).

